# Detecting molecular basis of phenotypic convergence

**DOI:** 10.1101/137174

**Authors:** Olivier Chabrol, Manuela Royer-Carenzi, Pierre Pontarotti, Gilles Didier

## Abstract

Convergence is the process by which several species independently evolve similar traits. This evolutionary process is not only strongly related to fundamental questions such as the predictability of evolution and the role of adaptation, its study also may provide new insights about genes involved in the convergent character. We focus on this latter question and aim to detect molecular basis of a given phenotypic convergence. After pointing out a number of concerns about detection methods based on ancestral reconstruction, we propose a novel approach combining an original measure of the extent to which a site supports a phenotypic convergence, with a statistical framework for selecting genes from the measure of their sites. First, our measure of “convergence level” outperforms two previous ones in distinguishing simulated convergent sites from non-convergent ones. Second, by applying our detection approach to the well-studied case of convergent echolocation between dolphins and bats, we identified a set of genes which is very significantly annotated with audition-related GO-terms. This result constitutes an indirect evidence that genes involved in a phenotypic convergence can be identified with a genome-wide approach, a point which was highly debated, notably in the echolocation case (the latest articles published on this topic were quite pessimistic). Our approach opens the way to systematic studies of numerous examples of convergent evolution in order to link (convergent) phenotype to genotype.

## 1 Introduction

Evolutionary convergence, which is strongly related to what is called homoplasy in cladistic, is a key concept in evolutionary biology (Stayton, 2015b; Pontarotti and Hue, 2016). The fact that several species, phylogenetically distant (e.g. placental and marsupial mammals), may share very similar traits cannot be explained without a certain amount of evolutionary convergence. For instance, Mahler *et al.* (2013) provide strong evidence of evolutionary convergence by carrying up a systematic study of various morphological characters of anoles having radiated onto four Caribbean islands. Convergence is indeed an important and quite common evolutionary mechanism, which concerns all kinds of traits: behavioral, morphological, developmental, molecular etc. (Losos *et al.*, 1998; Losos, 2009; Gallant *et al.*, 2014; Pfenning *et al.*, 2014; Vidal-García and Keogh, 2015; Ujvari *et al.*, 2015; Friedman *et al.*, 2016; Davis *et al.*, 2016). Widely studied, convergence has strong bearing on fundamental questions in evolution such as its predictability or the role of adaptation. A more practical motivation for studying this process, pointed out by Stayton (2015b), is that convergence is the only way to observe replicates of evolutionary events that cannot be repeated in controlled experiments for obvious reasons. Testing evolutionary hypothesis thus often requires to identify convergence events (Wake *et al.*, 2011).

As a key concept, convergence has been considered from various points of view, sometimes slightly different from one another. Intuitively, the main underlying idea is that there is convergence as soon as two or more species evolve independently similar traits. “Independently” stands here for the opposite of being derived from a common ancestral trait. In order to avoid confusion, let us start by giving a formal “working” definition of convergence, which will be discussed and refined in Section 2.2. We say that a molecular or phenotypic trait is *convergent* over two species s_1_ and s_2_ if the two following assertions are true:

1. both species s_1_ and s_2_ have the trait;
2. the MRCA of s_1_ and s_2_ does not have the trait.

Convergence may involve more than two species, say s_1_, s_2_, … and s_*n*_, all sharing a same trait. Note that, in this case, the trait can be said convergent as soon as at least a pair of species s_*i*_ and s_*j*_ have a MRCA which does not have the trait. The key point of this definition is that the trait has to evolve separately several times toward a same (or a similar) state different from the ancestral one(s). Some authors consider more precise definitions of convergence, by taking into account whether the ancestral states are different from one another (Zhang and Kumar, 1997) but we will not get into this level of detail here.

An important point is that deciding whether a given trait is convergent over species s_1_ and s_2_, requires to know whether their MRCA has this trait (actually, it may also require to quantify to what extent a trait is similar between two species but we will not consider this part of the problem). Since there is generally no definitive evidence about ancestral traits, convergence is always established with at least as much uncertainty as there is for ancestral reconstruction. Identifying convergent events is an important and difficult question in its own right. Several approaches have been developed for studying it (Revell *et al.*, 2007; Ingram and Mahler, 2013; Arbuckle *et al.*, 2014; Arbuckle and Speed, 2016; Stayton, 2015a; Speed and Arbuckle, 2017).

The question that we shall address here is slightly different from identifying convergence events. Given a binary character (typically the presence or absence of some phenotypic trait), which is assumed to be convergent for at least two extant species, we aim to detect genes showing molecular convergences, say at the amino acid level, which support, or at least are strongly consistent with, the convergence of the character. In particular, this is not the same as detecting molecular convergences *per se*, i.e. not related with a phenotypic character as considered in Zhang and Kumar (1997) and Storz (2016). To make it more concrete, the inputs of the question are:

- the phylogenetic tree of a set of extant species,
- the information about whether the phenotypic trait of interest is present for each extant species,
- the alignments of the clusters of orthologous genes of the extant species,

from which we aim to output a selection of genes which significantly support the convergence of the considered phenotype.

An expected outcome of identifying such genes is, first, a better understanding of evolutionary mechanisms leading to the acquisition of new traits, for instance with regards to an environmental pressure. Second, genes supporting the convergence are natural candidates for playing a role not only in the apparition of the phenotypic trait but also in its functions, possibly yielding new insights into the biological processes involved. This is thus an important question, which has been addressed by several previous works (Parker *et al.*, 2013; Foote *et al.*, 2015; Thomas and Hahn, 2015; Zou and Zhang, 2016). All the previous approaches have this in common that they first measure the strength of convergence, with regard to the character considered, for all sites of a given dataset and then select genes according to the convergence level of their sites. They do differ notably in the way of measuring the convergence level of sites.

A first class of measures of the convergence level of a site is conceptually very close to the definition above (Foote *et al.*, 2015; Thomas and Hahn, 2015; Zou and Zhang, 2016). Its main idea is to check if the species with the convergent trait show a same amino acid at the studied site and if this amino acid has been derived independently. To this end, the “convergent” amino acids are compared with the ancestral reconstructed ones, not necessarily at the MRCA level. For instance, Foote *et al.* (2015) compare the amino acid of each marine mammal with the reconstructed amino acid of its most recent ancestor having a terrestrial descendant. Assuming that the amino acids are accurately reconstructed allows counting the number of times that a given amino acid has been derived independently toward a species with the trait of interest (we shall see in Section 2.2 that this is not completely true). Since our approach was mainly designed to address some of its concerns, we will discuss ancestral reconstruction further in the next section. Let us just say that this kind of approaches heavily depends on the method used for reconstructing ancestral amino acids.

Another way of measuring the convergence level, which is used by Parker *et al.* (2013), consists in testing, for each site, the “real” phylogeny against an alternative phylogeny that separates the extant species having the convergent trait from the other ones. The convergence strength of a site is then measured in terms of ΔSSLS that is the difference between the log-likelihoods obtained from these two phylogenies. The approach is conceptually far from the definition of convergence, in the sense that ΔSSLS tests an evolutionary hypothesis corresponding to the alternative phylogeny, which is not obvious to interpret (Zou and Zhang, 2015b), rather than several independent mutations towards a same amino acid. We refer to Zou and Zhang (2015b) and Thomas and Hahn (2015) for a thorough discussion and a critical evaluation of this method.

The stage in which these approaches select genes from the convergence level of their sites is generally straightforward. For instance, Thomas and Hahn (2015) and Foote *et al.* (2015) considered genes which contain at least a convergent site, according to the measure used. Parker *et al.* (2013) ranked genes according to the mean ΔSSLS of their sites and considered the top-ranked ones.

Genome-wide detection of molecular signature of convergence is an emergent area of research which is still controversial. The article of Parker *et al.* (2013), about echolocation, was followed by two responses: from Thomas and Hahn (2015) and Zou and Zhang (2015b), which conclude that there is “no genome-wide protein sequence convergence for echolocation“. We propose here a new measure of the convergence level of a site, called *convergence index*, altogether with a statistical framework for ranking and selecting significant genes with regard to the convergence level of their sites. By applying our detection approach to the dataset of Thomas and Hahn (2015), still about echolocation, we draw the opposite conclusion to that of these authors. The set of genes significantly convergent between dolphin and microbat, the two echolocating species, show a very significant enrichment in GO-terms associated with audition in constrast to those detected from the other pairs of species in the dataset. These results provide an indirect evidence that molecular signatures of a phenotypic convergence may be detected with a suitable approach.

Source code of the software implementing the detection of molecular signatures of convergence is available at https://github.com/gilles-didier/Convergence.

## 2 Detecting genes supporting a phenotypic convergence

Given a set of extant species, among which some of them have a trait assumed convergent, the phylogenetic tree and alignments of orthologous genes of the extant species, the question is to identify the genes that show molecular convergences consistent with that of the trait. To this end, we follow the same general outline as the previous approaches, i.e. by first considering a convergence measure on alignment sites, then by selecting genes from the convergence level of their sites.

The ideas underlying the convergence measure that we developed, are conceptually close to the intuitive definition of convergence, thus to measures based on ancestral reconstruction. In fact, our measure, called *convergence index*, is essentially an attempt to address some concerns raised by ancestral reconstruction which are discussed in Section 2.1. The convergence index itself is presented in Section 2.2 (see also Appendix A).

The convergence index of a site is not directly used for measuring the strength of its convergence. We rather consider its significance under a null “neutral” evolution model in order to normalize effects due to the number of convergent extant species, to the phylogenetic tree and to the evolution rate of its gene (Section 2.3).

The last stage consists in selecting the genes which contain a significant number of sites detected convergent with regard to their index (Section 2.4).

### 2.1 Ancestral reconstruction approaches

Methods for identifying molecular signatures of convergence from ancestral sequences reconstruction (Foote *et al.*, 2015; Thomas and Hahn, 2015) raise several concerns, among which:

1. in order to decide whether there is convergence for a given site, one has to choose the ancestor nodes of which the reconstructed amino acids will be compared with those of convergent extant species;
2. ancestral reconstruction always comes with a certain amount of uncertainty, which is not taken into account by standard ancestral reconstructions;
3. approaches based on ancestral reconstruction implicitly assume that if one observes a same amino acid both at an ancestral species and at its direct descendant, then it was continuously present all along the branch (i.e. no mutation occurred during the corresponding time).

The first concern is not a big issue in the case where only two extant taxa have the convergent trait since, in this case, the MRCA is quite a natural choice. Things get more complicated when one has to deal with datasets containing a greater number of convergent extant species. Ancestral nodes to be compared with, have then to be chosen with regard to assumptions about whether they have the trait of interest. For instance, Foote *et al.* (2015) compare the amino acids of marine mammals with those reconstructed at their most recent ancestors having at least a terrestrial descendant, implicitly assuming that these ancestors were themselves terrestrial. Though there is strong evidence that assumptions made in Foote *et al.* (2015) make sense, reconstructing the evolutionary history of phenotypic traits is often much more challenging, and comes generally with a lot of uncertainty (Royer-Carenzi *et al.*, 2013; Royer-Carenzi and Didier, 2016). Moreover, results obtained this way heavily depend on the positions of selected ancestors in the tree, thus on the phylogenetic definition of the dataset.

The second concern may be easier to address. Ancestral reconstruction approaches based on stochastic evolution models are able to provide the probabilities of reconstructing one or another amino acid at a given ancestral node. This makes it possible to compute the expected number of convergent events in the sense of ancestral reconstruction approaches. Namely, given the amino acids of the convergent extant species and the ancestor nodes to which they are compared, the expected number of convergences under continuous time Markov model of evolution, can be directly computed (Zou and Zhang, 2015a). In a similar way, Zhang and Kumar (1997) and Castoe *et al.* (2009) were interested in the expected number of convergences between all pairs of branches of a phylogenetic tree by considering the posterior probabilities of all ancestor amino acids.

Let us remark that the definition of convergence given in the introduction, which is essentially at the basis of ancestral reconstruction approaches, is not completely consistent with the intuitive idea that there is convergence as soon as a trait (or an amino acid) appeared independently. There may indeed be an independent mutation toward an amino acid *X* inside a branch, even in the case where *X* is present at both nodes beginning and ending the branch. Figure 1 illustrates this point by displaying four different evolutionary histories of a site along a branch, all leading to observe *Alanine* (*A*) both at its beginning and at its end. All the histories but the one at the top-left of the figure, show an independent mutation toward *Alanine*, which will not be considered as such in an ancestral reconstruction framework, even if it deals with the reconstruction uncertainty like Zhang and Kumar (1997); Castoe *et al.* (2009); Zou and Zhang (2015a). In short, assuming that no mutation occurred on a branch which starts and ends with a same amino acid is an oversimplification which may lead to underestimate the actual number of molecular convergences.

**Figure 1:**
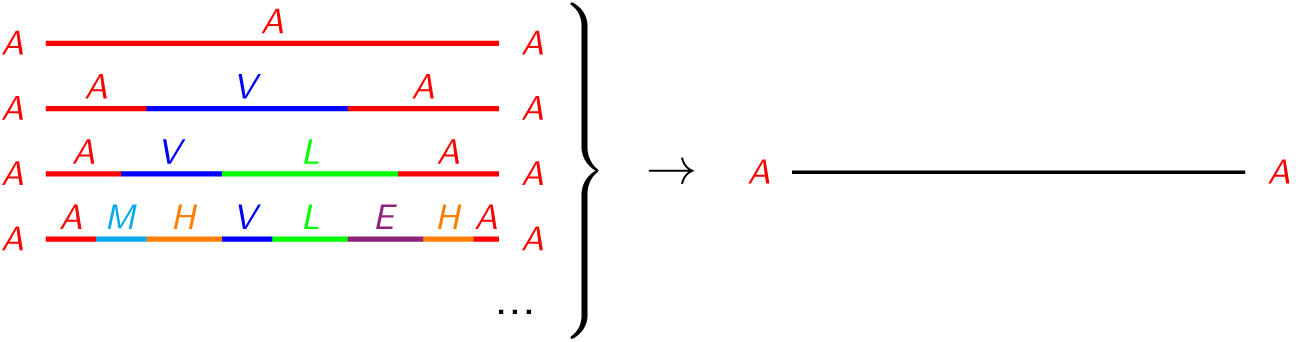
Several evolutionary histories (left) leading to observe amino acid *A* at the beginning and the end of a branch of a phylogenetic tree (right). The parts in red (resp. in blue, in green, …) correspond to the times when amino acid *A* (resp. *V*, *L*, …) was continuously present at the corresponding site.

### 2.2 Convergence index of an alignment site

In order to introduce our convergence measure, let us start by assuming that the whole evolutionary history of a site is known. By whole evolutionary history, we mean that the amino acid present at all lineages and at all times encompassed by the phylogenetic tree is known (i.e. we know which amino acid is present not only at the nodes but anywhere in the tree, including inside branches). In this situation, and for all amino acids *X* present at a convergent extant species, it is straightforward to count the number of mutations towards *X* which are conserved until an extant species with the convergent trait. This number reflects intuitively the extent to which mutations toward *X* supports the convergence of the trait for the site considered. It will be referred to as the *number of independent apparitions of X*. Figure 2 displays several evolutionary histories leading to different number of independent apparitions of amino acid *A*, which correspond to the number of starting points of the red parts of branches in the figure. Note that it may occur that the amino acid considered (*A* in Figure2) is not present at all the extant species having the convergent trait (e.g. evolutionary history at the bottom-right of Figure 2).

**Figure 2:**
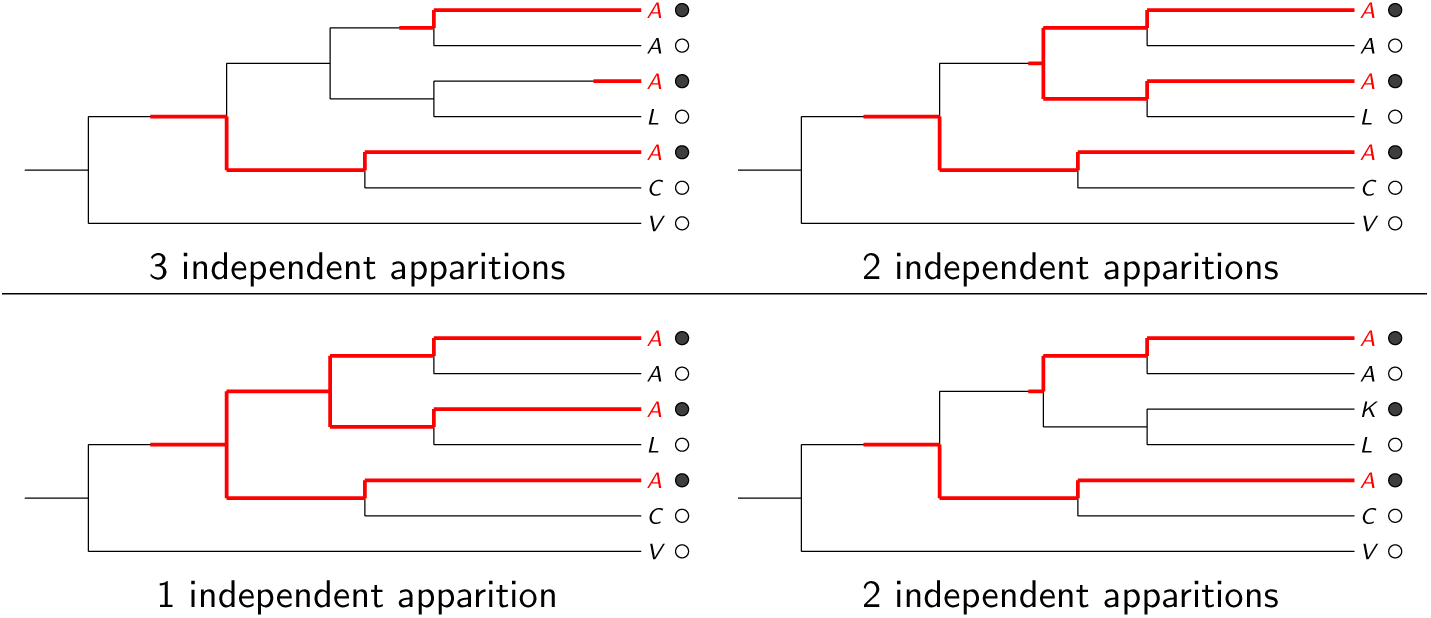
Independent apparitions of amino acid A with regard to a given phenotypic trait assumed convergent. Extant species are represented with • or ∘ depending on whether they have the convergent trait. Red parts of branches end at all tips with both the convergent trait and amino acid A, and keep go back in time as long as A was continuously present at the site.

Unfortunately, in a real situation, we don‘t have access to the whole evolutionary history of a site, but only to the amino acids of the extant taxa. All is not lost, however, since we are able to compute the expected number of independent apparitions under a standard continuous time Markov model of evolution (Yang, 1994). We present in Appendix A, a method for computing the expected number of independent apparitions of a given amino acid with regard to a phylogenetic tree, the traits of extant species and an alignment site, under a continuous time Markov model of evolution.

The convergent index of a site is defined as the maximum of the expected numbers of apparitions over all amino acids present at convergent extant species, conditioned on amino acids of all the extant species (i.e. the corresponding alignment column, see Equation 1 of Appendix A).

Let us note that the concerns stated at the beginning of Section 2.1, do no apply to the convergence index, since:

1. computing the expected number of apparitions of an amino acid does not require to select any ancestor node;
2. it does take into account the uncertainty due to the stochastic nature of evolution, since it is an expectation under a probabilistic model;
3. our calculus distinguish between the case where there is no mutation all along a branch of the phylogenetic tree and the case where an ancestor and its direct descendant share a same amino acid (Appendix A).

The computation of the convergence index requires a continuous time Markov model of sequence evolution, which is generally given by its substitution rate matrix (Whelan and Goldman, 2001). In order to compute likelihoods over phylogenetic trees, this matrix has to be multiplied by a constant rate, standing for the evolution speed with regards to the time unit of the branch lengths. The choice of this rate has a great influence on the convergence index. In particular, a high rate leads systematically to convergence indexes almost equal to the number of convergent species. After trying several alternatives, we devised a heuristic for calibrating the evolution rate used for computing the convergence index from the phylogenetic tree and the convergent species, which ensures that the convergence index takes values over a range as wide as possible (Section 5.3).

### 2.3 Signicance of a site

By construction and for all amino acids *X*, the expected number of independent apparitions of *X* ranges between 0 and the total number of convergent extant species. It follows that the convergence index heavily depends on the number of convergent extant species. Another factor which influences the expected number of apparitions of an amino acid is the evolutionary rate of the site. For instance, no molecular convergence can be identi ed from a completely conserved site. Figure 3 displays the simulated distributions of the convergence index over two different trees, having respectively three and two convergent species, and at evolutionary rates 10 and 30. We do observe that these distributions are quite different between each other. In particular, distributions simulated with rate 10 do not have the same general shape as those simulated with rate 30. Note that the rate used for computing the convergence index is here fixed, constant over all the plots, and different from those used for the simulations. Convergence indexes of the bottom row distributions are bounded by 2 while those of top row distributions may actually go until 3, the number of convergent species, though the corresponding probabilities are very low and (at most) barely noticeable on the plots (Figure 3).

**Figure 3:**
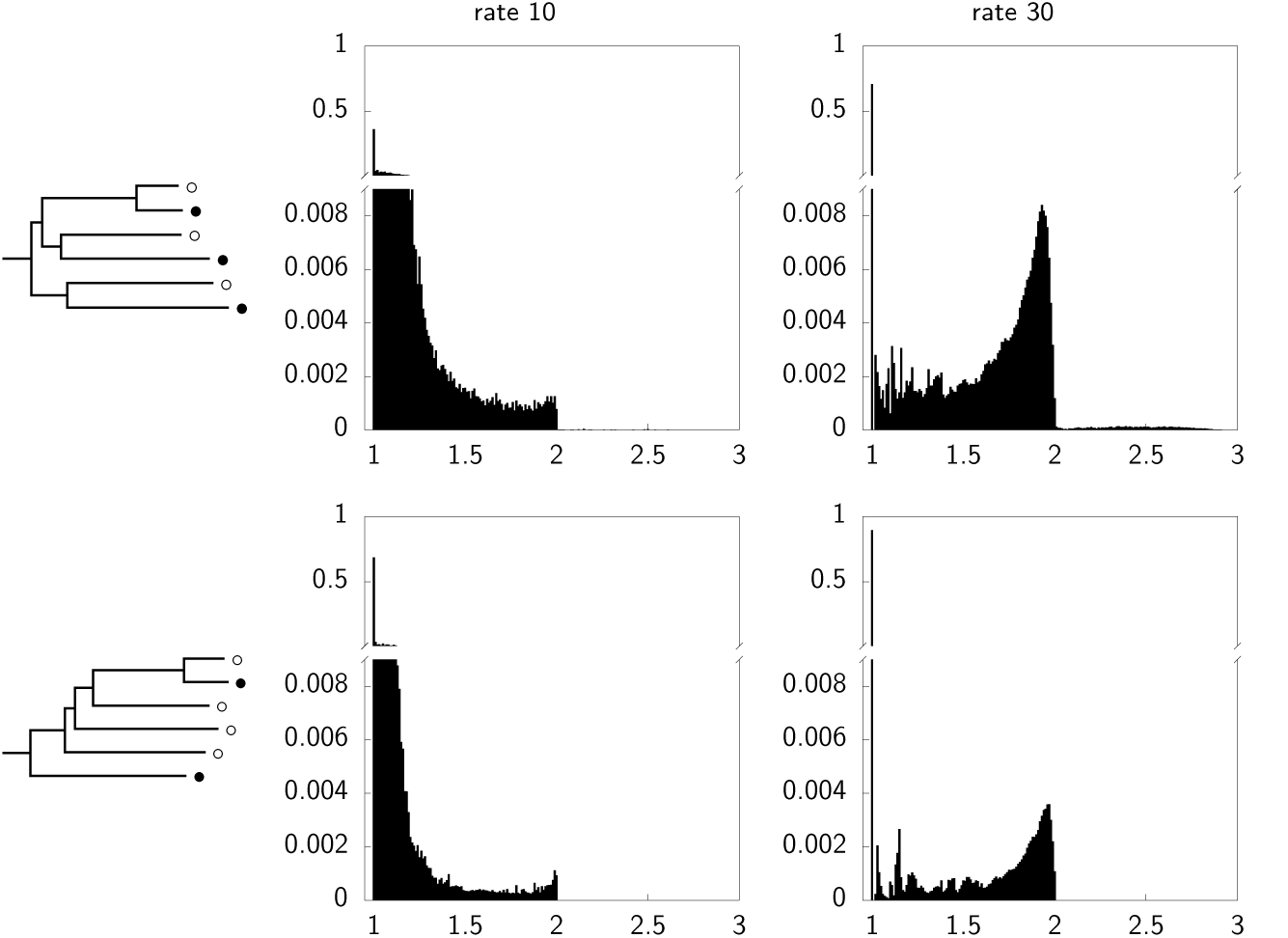
Simulated distributions of the convergence index under neutral evolution. Distributions of each line are simulated from the tree displayed in Column 1 at evolution rates 10 (Column 2) and 30 (Column 3). The tree of the top row (resp. of the bottom row) has three (resp. two) convergent species (represented with •).

In order to decide whether a site is convergent, we thus have to normalize its convergence index with regard to its evolution rate, the phylogenetic tree, and the number and positions of extant species with the convergent trait. To this end, we consider the *p*-value associated to its convergence index with regard to the empirical distribution of convergent indexes computed from the same phylogenetic tree and the same set of convergent extant species but for sites obtained by simulating neutral evolution of an amino acid on the phylogenetic tree, under an evolution model of which parameters are estimated from the whole gene to which the tested site belongs. We insist on the fact that the model used for simulating sites does not have to be the same as the one used for computing the convergence index. Convergence index is treated here as a statistics of the site of which we evaluate the distribution under an evolution model of its gene (this model may take into account rates heterogeneity etc.). In the current implementation of the method, the same substitution rate matrix is used both for convergence indexes and for simulations but convergence indexes are computed with a single evolution rate while simulations are performed from evolution rates drawn from a discretized Gamma distribution. Computing convergence indexes from the exact same model as for simulating worked as well but was several times more time-consuming.

#### 2.4 Significance of a gene

In our context, assessing the significance of a gene requires to combine the (empirical) *p*-values of its sites. Since combining *p*-values is a question of broad interest, several methods have been developed for performing this task (Loughin, 2004). The widely used “quantile” approaches such as Fisher and truncated product are not well suited to our particular question. Empirical *p*-values are prone to uncertainty, notably for the smallest ones which have the greatest influence on these methods. We rather follow Wilkinson (1951) and start by choosing a significance level *γ*. A site is said *convergent at a significance level γ*, or *γ -convergent*, if the probability of observing a convergence index greater of equal to its own convergence index is smaller or equal to *γ*, in the empirical distribution associated to its gene as described in Section 2.3. We developed an adaptive sampling scheme which determines the number of simulations required for ensuring a given confidence level to the number of *γ* -convergent sites of a gene, with regard to its length and *γ*. All genes are next associated with the number of *γ* -convergent sites that they contain. By assuming independence between sites, the number of convergent sites of a gene of length *L* follows a binomial distribution of parameters (*γ, L*). The (combined) *p*-value of a gene,that is the probability of observing a number greater or equal than the observed number of convergent sites in this binomial distribution, is thus straightforward to compute. This *p*-value has to be corrected for multiple testing with regard to all the genes/alignments in the dataset, in order to give the final significance of this gene.

## 3 Results

### 3.1 Comparison of 3 measures of convergence of a site

In order to assess the accuracy of measures of convergence level, we simulated the evolution of non-convergent and convergent sites on the tree of Thomas and Hahn (2015) (Figure 4-left-top). We compared 3 measures, namely the convergence index, the number of convergence observed from the ancestral reconstruction (Foote *et al.*, 2015) and the ΔSSLS (Parker *et al.*, 2013). We used the tree of Thomas and Hahn (2015) with the same convergent extant species, also displayed in Figure 5. The ΔSSLS measure was computed with an alternative tree built following the ideas of Hypothesis H2 in Parker *et al.* (2013) (Figure 4-left-bottom). Following Foote *et al.* (2015), we compare the amino acids of convergent extant species with those reconstructed at their most recent ancestors with least a descendant without the convergent trait, in order to count the number of convergent events for the ancestral reconstruction method. Since all the sites were simulated on the same tree with the same convergent species and under a same evolution rate, it is not required to normalize the convergence indexes with regard to their empirical distribution. They are thus used directly.

**Figure 4:**
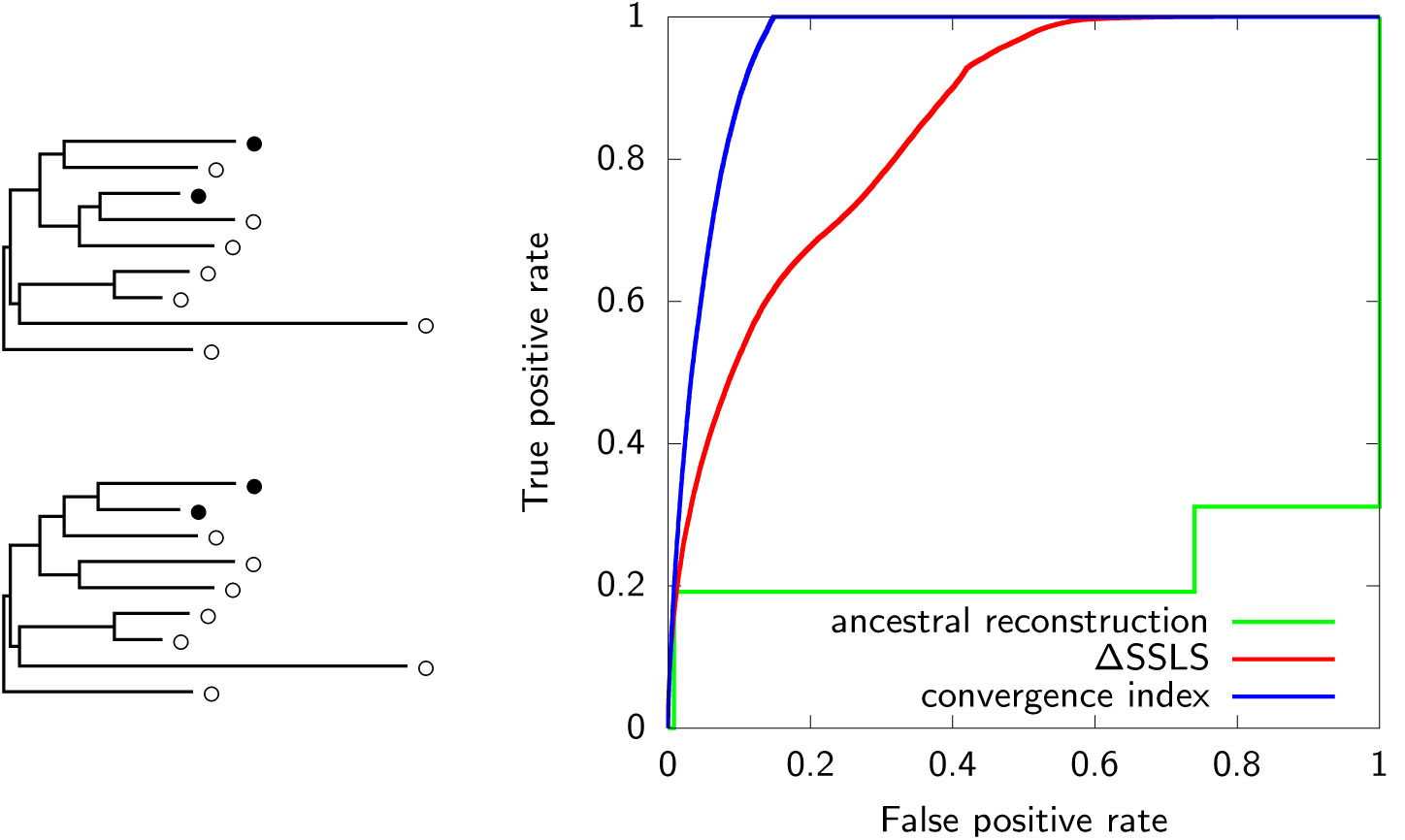
(left-top) Tree used for simulating non-convergent and convergent protein site. (left-bottom) Alternative tree used for SSLS computation. Extant species are represented with • or ∘ depending on whether they have the convergent trait. (right) RoC curves obtained from 100, 000 simulations. The closer a RoC curve is to the upper left corner, the higher the accuracy of the corresponding convergence measure.

**Figure 5:**
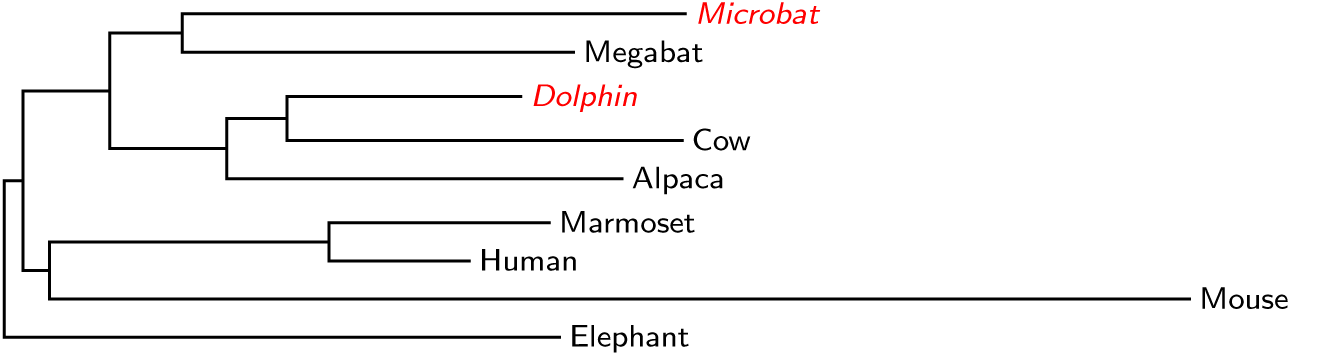
Phylogenetic tree from Thomas and Hahn (2015). Among the nine extant species, dolphin and microbat (in red italic) have echolocation abilities.

The relevance of the convergence measures was next assessed with regard to their ability to distinguish between the simulated non-convergent and convergent sites (the status of all simulated sites is known). Figure 4-right displays the results obtained for the ancestral reconstruction, ΔSSLS and the convergence index. The RoC curves (Zhou *et al.*, 2009) reporting the results of each measure, show that the convergence index better discriminates between convergent and non-convergent sites than the two other measures (Figure 4-Right). In this particular example, the convergence index identifies 99% of the convergent sites with an error rate of 1.4%, with a suitable threshold.

### 3.2 Convergent genes related to echolocation

We applied the approach described in Section 2, to the dataset of Thomas and Hahn (2015). This dataset was designed for studying the apparition(s) of echolocation abilities in mammals, like that of Parker *et al.* (2013). It contains 6,332 alignments of orthologous genes from 9 mammal species, among which two have echolocation abilities (dolphin and microbat), and the phylogenetic tree of these 9 species (Figure 5 and Section 5.6).

Convergent sites are detected at a confidence level of 10^−4^ according to the empirical distribution simulated with regards to the genes to which they belong (Sections 5.7). We next computed the (binomial) *p*-values of all genes with regard to the number of convergent sites that they contain, ranked the genes according to their *p*-values and performed a Benjamini-Hochberg correction for multiple-testing (Section 5.8). We finally selected the genes with corrected *p-* values smaller than 5 *×* 10^−2^, that give us the set of convergent genes. The detection of convergent genes was performed between all pairs of species in the dataset for control purposes.

Figure 6 displays the number of genes detected convergent for all pairs of species. No convergent genes were found for pairs of sister species (cow-dolphin, microbat-megabat and human-marmoset). But there is a certain amount of convergent genes for almost all the other pairs of species, ranging from 23 to 152 genes, except for the pair human-mouse which has only 2 convergent genes. We did not observe more convergent genes between the two echolocating species than between the other pairs, which is consistent with what was observed by Thomas and Hahn (2015).

**Figure 6:**
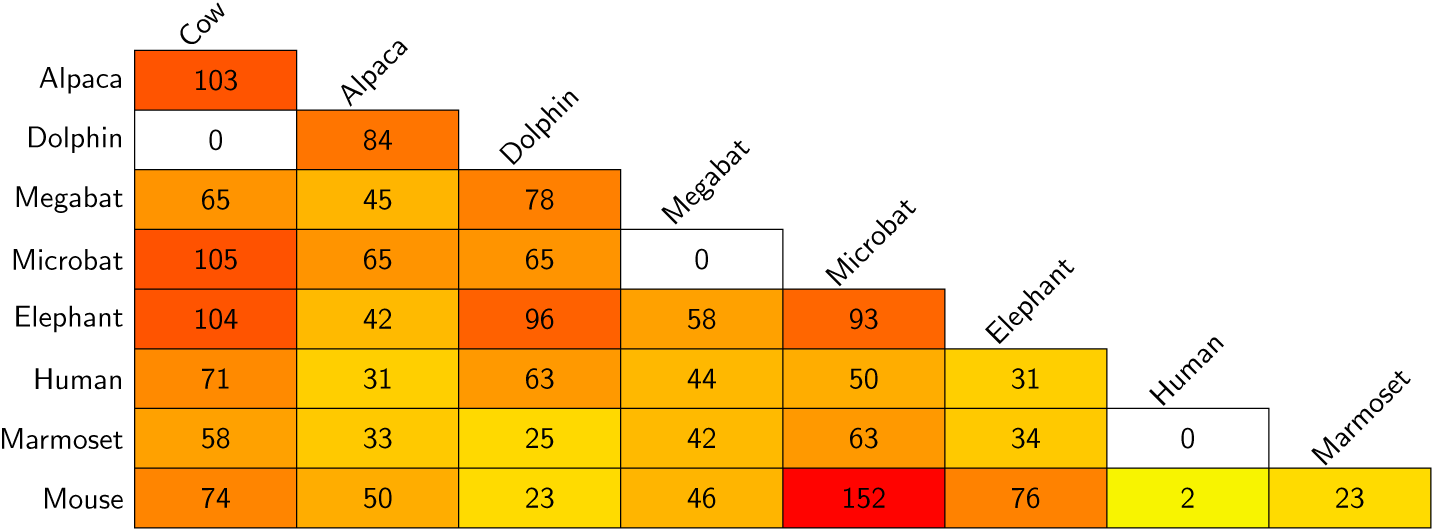
Number of genes detected convergent for all pairs of species of the dataset.

In order to assess if the convergent genes detected between dolphin and microbat were related to echolocation, we tested their enrichment with regards to GO-terms involved in audition (see Section 5.9). We performed the same test for all pairs of species, in order to ensure that there is no bias leading to observe more convergence on genes associated with these particular GO-terms. Results are displayed in Figure 7, which shows that the set of convergent genes between dolphin and microbat is by far the most significantly enriched in audition-related annotations. The corresponding Fisher‘s exact test *p*-value, i.e. 6.66 *×* 10^−5^, is (at least) three orders of magnitude lower than those of other pairs of species.

**Figure 7:**
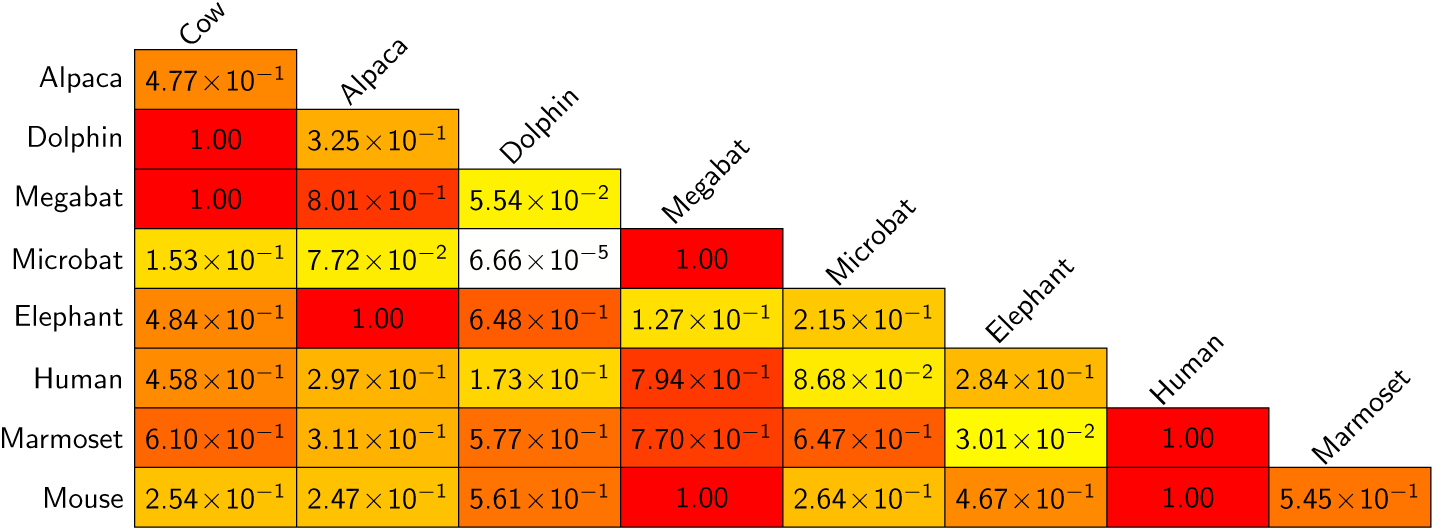
Enrichment *p*-values, with regard to the “audition-related” GO-terms, of the genes detected convergent for all pairs of species of the dataset.

Note that Prestin (a.k.a. SLC26A5), known to be involved in echolocation (Li *et al.*, 2010; Liu *et al.*, 2010, 2014), is the 3^rd^ most significant convergent gene out of the 6, 332 ones of the dataset. Still for studying echolocation, Shen *et al.* (2012) screened three genes, namely CDH23, PCDH15 and oToF, pointing out that “Convergent evolution and expression patterns of oToF suggest the potential role of nerve and brain in echolocation“. The two first genes were not in the dataset of Thomas and Hahn (2015) but otoferlin (oToF) was well detected convergent with our approach (at the 54^th^ rank). Davies *et al.* (2012) found signatures of sequence convergence in TMC1 and DFNB9 (aka PJVK). These genes are respectively the 5^th^ and 11^th^ most significantly convergent with our method (Supplementary Information). Among the seven genes pointed out by Parker *et al.* (2013) as previously reported for showing convergence and/or adaptation in echolocation, four were present in the dataset. We detected all of them (in bold in Table 1 of Supplementary Information). Table 1 displays the significant GO annotations of genes convergent for the echolocating pair, that are related with audition (as defined in Section 5.9) and their Fisher‘s exact test *p*-values without multiple testing correction with regard to the total number of GO-terms.

**Table 1:** Significant audition-related GO annotations of genes detected convergent between dolphin and microbat.

| Fisher’s exact *p*-value | GO ID | Description | Genes |
| --- | --- | --- | --- |
| $4.36 \times 10^{-4}$ | GO:0007605 | sensory perception of sound | SLC26A5, TMC1, DFNB59, COL11A1, OTOF |
| $4.59 \times 10^{-3}$ | GO:0050910 | detection of mechanical stimulus involved in sensory perception of sound | TMC1, COL11A1 |
| $7.79 \times 10^{-3}$ | GO:0090102 | cochlea development | SLC26A5, OTOF |
| $1.89 \times 10^{-2}$ | GO:0007420 | brain development | PTPRZ1, RELN, MED1, KCNAB1 |
| $3.10 \times 10^{-2}$ | GO:0060117 | auditory receptor cell development | TMC1 |

## 4 Discussion

There is mounting evidence that phenotypic convergence has detectable molecular basis for many phenotypic characters, including echolocation (Li *et al.*, 2010; Liu *et al.*, 2010). Despite this fact and the importance of this matter, there is still no consensus method for the genome-wide identification of the molecular signatures of a given phenotypic convergence. We presented here, first, a new measure for evaluating the extent to which an alignment site supports a phenotypic convergence and, second, a statistical framework for detecting significant genes from the convergence level of their sites.

A first result is that our convergence measure has better performance for detecting convergence than two previous measures on simulated sites. In particular, the convergence measure based on ancestral reconstruction (the “historical” approach) showed poor results on discriminating between convergent and non-convergent simulated sites. This is not a surprise in view of the concerns we listed in Section 2.1 and of the lack of definition of this approach, which basically decides if the amino acids of the convergent species are convergent or not, without considering any nuance between the two situations. The ΔSSLS approach has better results than the ancestral one but is outperformed by the convergence index whatever the alternative tree we tried.

Our detection pipeline did not return a greater amount of convergent genes between the two echolocators than between the other pairs of species. This point was not completely unexpected since dolphin and microbat evolve in very different environments and, at first glance, do not share more phenotypic characters than the other pairs (echolocation excepted). Nevertheless, the fact that the amount of convergent sites or genes detected between echolocators was not greater than between other pairs was put forward as an argument against the detectability of the molecular basis of echolocation (Thomas and Hahn, 2015; Zou and Zhang, 2015b). We argue that this point is not conclusive since it is based on the assumption that non-echolocating pairs have no phenotypic convergence (Thomas and Hahn (2015) used them for determining a null distribution). The actual amount of phenotypic and molecular convergence between species remains difficult to predict since it may involve “traits” not obvious to observe (e.g. metabolic pathways, proteins binding etc.). Evaluating the actual extent of convergence between species needs further investigations which are out of the scope of the present work. At this point, Figure 6 suggests either that convergence is quite a common mechanism of which molecular traces are detectable, or a possible issue in the approach.

The relevance of the results obtained with our pipeline was assessed with regard to the particular phenotype studied in the dataset. Since echolocation requires special hearing capacities, one expects genes detected convergent between the two echolocating species to be, for at least some of them, related to audition. This point is clearly observed in Figure 7 (see also the Supplementary Information). on the contrary, Thomas and Hahn (2015) found no evidence of sensory enrichment in genes detected convergent with the ancestral reconstruction approach. Though Parker *et al.* (2013) observed several hearing genes among the top 5% with the highest ΔSSLS, they did not provide any statistical support for this point (they obtained 117 genes among which only 4 were also detected convergent by our method). They showed that a selection of hearing (and sensory) genes have ΔSSLS higher than expected but not at a level extremely significant and without checking the non-echolocating pairs as pointed out by Thomas and Hahn (2015).

The significance threshold, γ which is used for deciding if a site is convergent with regard to the empirical distribution associated to its gene, is a crucial parameter of our approach. Its choice is up to the user and relies on what is expected about molecular convergence, i.e. a signal diffuse or concentrated on a few sites. We tested several values γ of from 10^−3^ to 10^−5^. For all cases but *γ* =10^−3^, *p*-values of the “audition enrichment” of the set of genes detected between the echolocating pair were at least one order of magnitude lower than those of the other pairs (Supplementary material). Though this is not an absolute rule, the number of detected genes tends to decrease with γ for all pairs of species. Significance with regards to audition-related GO-terms peaks at *γ* = 5 *×* 10^−5^ for the echolocators but only 36 convergent genes are detected at this level. There are only 7 genes left for *γ* = 10^−5^ and there is no point in considering lower thresholds.

Though we aim to provide a rigorous framework for detecting convergent genes, the relevance of our results heavily depends on our assumptions with regard to protein evolution. Since deciding if a site is convergent relies on simulations from the evolution model chosen, the more realistic this model, the more accurate the results we get. By “evolution model“, we mean here both the modeling of amino acid substitutions and that of the rate heterogeneity along a gene. The current version of the detection pipeline is based on the widely used model WAG+discretized Gamma distribution as implemented in PAML (Yang, 1994). Any evolution model, whatever its sophistication, may be easily plugged into the detection pipeline, since it is only used for simulating empirical null distributions. We plan to test more realistic models in the future.

Though there is still room for improvement, the fact that genes we detected convergent for the echolocating pair are annotated with audition-related GO-terms at a significance level far greater than for the other pairs of species, constitutes a proof of concept that a genome-wide approach may identify molecular basis of a given phenotypic convergence. Whether such an identification was possible was debated, notably in a genome-wide context. First, detecting convergent genes is possible only if at least some of the mutations leading to the phenotype involve the same sites, thus the same genes, which corresponds to strong constraints on evolution. Second, the preceding condition is not sufficient to ensure that convergent sites are detectable. In a genome-wide context, this also requires a rigorous statistical framework for evaluating the significance of the molecular convergences observed at sites, in other words, a way of distinguishing convergence signal from evolutionary noise. This latter point is not a real concern when genes in which molecular signatures are expected are known *a priori* (Zhang, 2006; Ujvari *et al.*, 2015) but is essential for dealing with thousands of genes.

Since Conte *et al.* (2012) estimated that phenotypic convergence involves the same genes in between a third and a half of the cases, the numerous occurrences of phenotypic convergence observed in nature constitutes a huge dataset that can be used for studying relations between genotype to phenotype.

## 5 Material and Methods

### 5.1 Detection pipeline overview

The detection pipeline is schematically displayed in Figure 8. In order to detect molecular signatures of a given convergent character inside a set of genes, it takes as input the alignments of orthologous sequences of the genes for a set of species containing the convergent ones, the phylogenetic tree of these species and the information about whether they carry the convergent character. Users have to provide four parameters: a substitution matrix *M*, suited to the type of sequences considered, a significance threshold for deciding which sites are convergent and a significance threshold for deciding if a gene is convergent with regard to its length, the convergent sites that it contains and the total number of genes. The execution of the pipeline follows three stages. Stage 1, “Method calibration“, determines the evolutionary rate *µ* used for computing the convergence index with matrix *M* (Section 5.3). Stage 2 consists in treating all alignments/genes of the dataset by (i) estimating the parameter *α* of the discretized Gamma distribution for the evolutionary rates of the alignment protein from matrix *M* (Yang, 1994), (ii) simulating the empirical distribution of the convergence index from the estimated parameter *α* with *M*, (iii) computing the convergence index of all sites with the method rate *µ* under *M* and (iv) determining the *p*-values associated to alignments/genes with regard to parameter *γ*, the number of sites of significance smaller than *γ* and the length of alignments/genes. In Stage 3, *p*-values are corrected for multiple testing. Finally, alignments/genes with corrected *p*-values smaller than parameter *β* are output. The pipeline also returns the complete list of genes, sorted according to their *p*-values, and the positions of their *γ* -convergent sites (Supplementary information).

**Figure 8:**
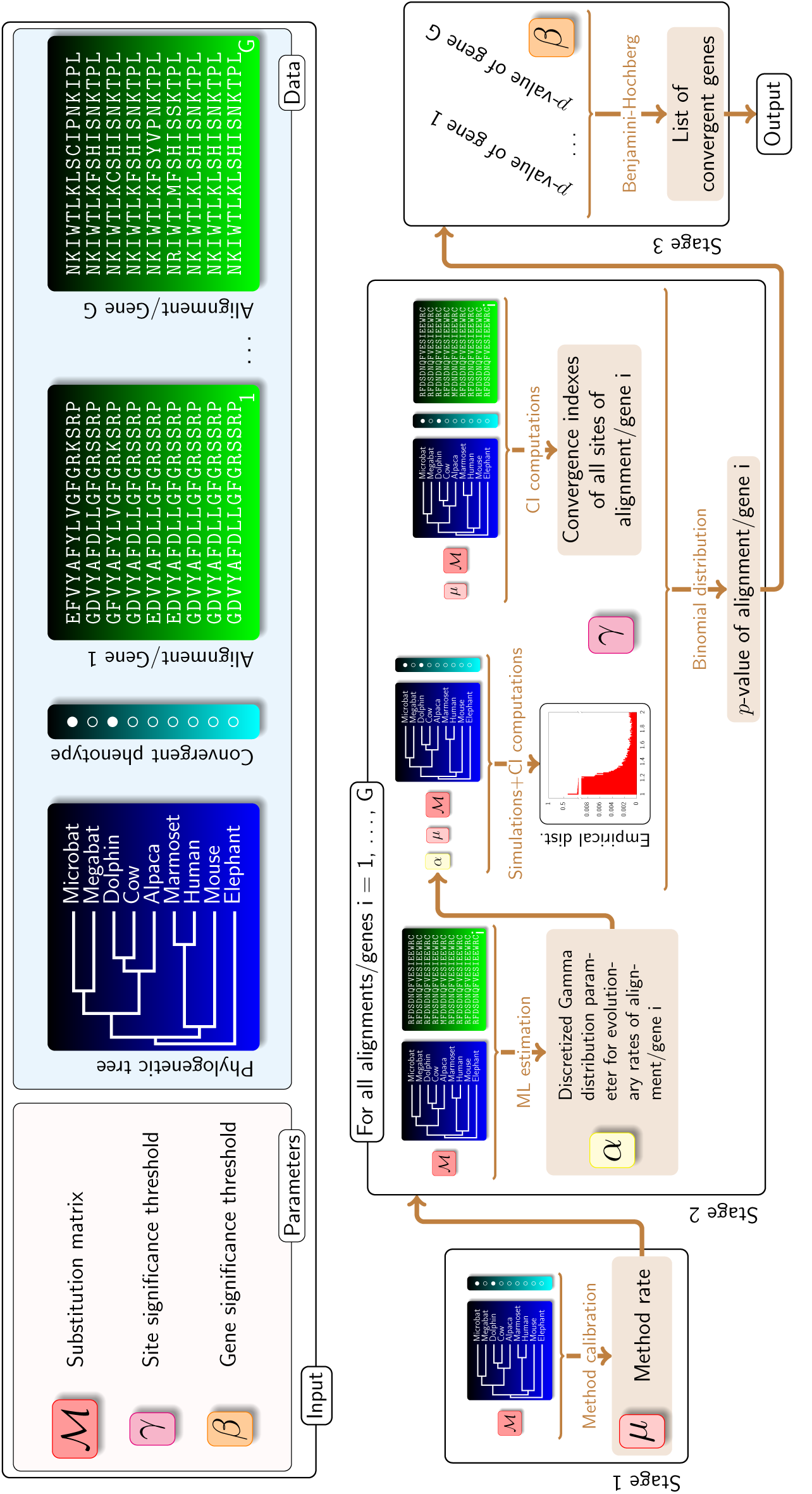
Schematic of the detection pipeline. *ML* stands for *Maximum Likeli-hood* and *CI* for *Convergence Index.*

### 5.2 Convergence index of a site

The convergence index of a site of a gene is determined from the expected number of independent apparitions of all amino acids as presented in Appendix A. These expected numbers are computed under a WAG model (Whelan and Goldman, 2001), with the evolutionary rate determined as presented in Section 5.3.

### 5.3 Method calibration

The method is calibrated with regard to the phylogenetic tree, the convergent characters and the evolution model. our calibration heuristic consists in finding the evolutionary rate (or more generally the evolution model parameters) which is such that the alignment column with amino acid *Leucine* at all entries, has a convergence index equal to 1.1. Note that the amino acid *Leucine* was arbitrarily chosen and so was the value 1.1. Calibrating the evolutionary rate of the method with another amino acid and/or any value slightly greater than 1 leads to similar results.

### 5.4 Simulating non-convergent sites

Simulated non-convergent sites, used for plotting the RoC curves of Figure 4-left-top, were obtained by simulating the evolution of an amino acid on the tree displayed in Figure 4, under the WAG model with an evolutionary rate of 10 (Whelan and Goldman, 2001) and by keeping only the amino acids of the extant species which give us our alignment columns.

### 5.5 Simulating convergent sites

Simulated convergent sites were obtained in two stages. First, we simulated the evolution of an amino acid in the very same way as for a non-convergent site. Second, for all sites such simulated, we randomly picked an extant species with the convergent trait and we set the amino acids of all the other convergent extant species, to that observed at the one picked. This way, we get an alignment column (i.e. a site) in which a same amino acid occurs at all the entries corresponding to the convergent species.

### 5.6 Biological dataset

We used the dataset of Thomas and Hahn (2015). It contains the phylogenetic tree displayed in Figure 5 and 6,400 protein alignments, among which we kept only the ones with *Ensembl* IDs which mapped to *RefSeq* IDs. This left 6, 332 alignments of length ranging from 1 to 4, 822, with an average length of approximately 295 amino acids.

### 5.7 Significance of a site

The empirical distributions used for evaluating the significance of convergence indexes of sites are obtained by simulating neutral “non-convergent” sites as described in Section 5.4, under the standard evolution model with multiple rates drawn from the 4-values discretized Gamma distribution estimated from the gene containing the site (Yang, 1994).

### 5.8 Significance of a gene

All sites with a convergence index significant at a level smaller or equal to γ = 10^−4^ with regard to the empirical distribution corresponding to its gene, are considered convergent. The total number of *γ*-convergent sites of a gene allows us to compute a *p*-value from a binomial distribution of parameters γ= 10^−4^ and the length of the gene. Finally, the *p*-values of genes are corrected for multiple-testing by using the Benjamini-Hochberg procedure (Benjamini and Hochberg, 1995).

### 5.9 GO-term enrichment

The *RefSeq* IDs were fetched from all *Ensembl* IDs of the dataset of Thomas and Hahn (2015) by using the perl API of *Ensembl* (see http://www.ensembl.org/ info/docs/api/core/core_tutorial.html). We next obtained all the GO-terms associated to each *RefSeq* ID *via QuickGO* (https://www.ebi.ac.uk/ QuickGO/WebServices.html). The set of GO-terms associated to audition is the union of GO-terms returned by searching the keywords “sound“, “hearing“, “auditory” and “cochlea” in the *AmiGO 2* site (http://amigo.geneontology.org/amigo/). Exact Fisher‘s tests were performed with the 6, 332 genes of the dataset as background.

## Authors contributions

OC contributed to the software development, fetched the data and ran the tests. MR-C provided statistical insights and a careful reading of the manuscript. PP originated the biological side of this work, provided evolutionary insights all along its development and most of the references. GD provided the original idea of the detection approach, supervised its development and its evaluation, derived the mathematical part, led the software development and wrote the manuscript. All authors read and approved the final manuscript.

## Acknowledgement

All authors wish to thank Gregg W.C. Thomas and Matthew W. Hahn for the quality of their dataset and for kindly providing it. GD thanks Bastien Boussau for many discussions about evolutionary convergence.

## A Convergence index of a site

Let *𝒯* be a phylogenetic tree. For all nodes *n* of *𝒯* (tips included), we put

- *𝒯* _*n*_ *for* the length of the branch leading to *n*,
- *𝒯* (*n*) for the subtree rooted at *n*,
- *𝒞* (*n*) for the set of direct descendants of *n*.

Let us put *ℒ* for the set of tips of *𝒯* and *𝒳* for the subset of tips which contains all the extant taxa having the convergent trait (*| 𝒳 |* designs the number of species in *𝒳*).

We present here the computation of the convergence index of a protein site. The exact same method can be applied on a site of a DNA or a codon sequence. It just requires to change the evolution model accordingly.

In the present context, a site is a column of a given alignment of orthologous proteins of the extant species of *𝒯*. For all species *ℓ* ∈ *ℒ*, we put *α*(*ℓ*) for the amino acid present at the site considered, i.e. the entry *ℓ* of the corresponding alignment column.

For a given amino acid *A* (here and thereafter, ‘*A*’ is a generic notation for designing an arbitrary amino acid and does not stand for ‘*Alanine*’), the *A-expectation* of the site is defined as the expected number of times that a mutation towards *A* is observed (from an amino acid different from *A*), after which *A* is continuously conserved until at least a tip of *𝒳* (see Section 2.2 and Figure 2). This expectation is computed under a standard continuous time Markov model of evolution (π*Q*), where *π* is the amino acid distribution at the root of the tree and *Q* is the infinitesimal generator of the continuous Markov chain modeling the mutation process (see for instance Felsenstein (1981)). A *continuous A-path* is a path of the phylogenetic tree starting at some point (possibly inside a branch) and ending at a tip having the convergent trait, in which the amino-acid *A* is continuously present (represented in red in Figure 2). We will say that a continuous *A*-path *passes through* a node *n* of the tree if it starts inside an ancestor branch of *n* and ends at a tip, which has the convergent trait and which descends from *n*. By construction, the *A*-expectation is equal to the expected number of starting points of continuous *A*-paths in *𝒯* (Figure 2).

In order to compute the *A*-expectation of a site, we define, for all nodes *n* of *𝒯* and all integers 0 *≤ k ≤ |*𝒳*|*,

- C_*A*_(*n, k*) as the probability of observing *k* starting points of continuous *A-*paths in the subtree rooted at *n* and a continuous *A*-path passing through *n*, which implies that *A* is the ancestral amino acid at *n*, conditioned on having amino acid *A* at node *n*;
- for all amino acids *X*, B_*A*_(*n, X, k*) as the probability of observing *k* starting points of continuous *A*-paths in the subtree rooted at *n* with no continuous *A*-path passing through *n*, conditioned on having amino acid *X* at node *n*.

We shall proceed in a very similar way as for computing the likelihood of the tree (Felsenstein, 1981). Before establishing recurrence formula determining the above conditional probabilities of an internal node from those of its children, we start with base cases, i.e. the conditional probabilities of tips. Since the subtree pending from a tip *n* is empty, both C_*A*_(*n, k*) and B_*A*_(*n, X, k*) are zero for all *k >* 0. If *n* is a tip having the phenotype, then there is a continuous *A*-path passing through it, if and only if amino acid *A* is present at *n*. Namely, it follows that we have for all integers 0 *≤ k ≤ |X |*,

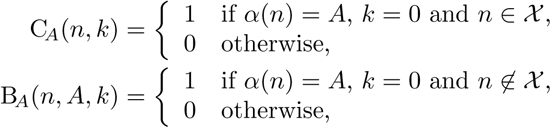

and, for all amino acids *Y ≠ A*,

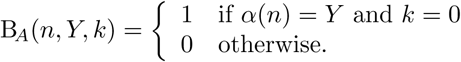

Let us put:

- p_C_(*A, t*) for the probability of keeping an amino acid *A* without mutation all along a branch during time *t* under the evolution model (*π, Q*), namely 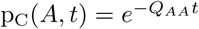 (e.g. top-left evolutionary history of Figure 1),
- p_M_(*X, Y, t*) for the probability of going from an amino acid *X* to an amino acid *Y* in time *t*, namely the entry (*X, Y*) of the transition matrix *e*^*Qt*^.

For all internal nodes *n* and all *c ϵ C*(*n*), we define the tree 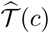 as the subpart of *𝒯* which contains the node *n*, the branch going from *n* to *c* and the subtree *𝒯* (*c*) (i.e. the subtree rooted at *n* in which all the descendants of *n* other than those in the subtree *𝒯* (*c*) were pruned). Note that 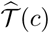 is rooted at *n*, the direct ancestor of *c*, and not at *c* itself.

We then define

- 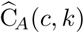 as the probability of observing *k* starting points of continuous *A*-paths in 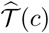 and a continuous *A*-path passing through *n*, conditioned on observing the amino acid *A* at *n*;
- 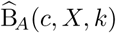 as the probability of observing *k* starting points of continuous *A*-paths in 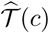 and no continuous *A*-path passing through *n*, conditioned on observing the amino acid *X* at node *n*.

Let us compute the conditional probabilities 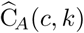 and 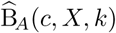 from the conditional probabilities C_*A*_(*c, ℓ*) and B_*A*_(*c, X, ℓ*) for all amino acids *X* and all integers 0 *≤ k ≤ |*𝒳*|*.

If there is a continuous *A*-path passing through *n* in 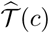, it necessarily passes through the node *c* and it is not interrupted in the branch between *n* and *c*. There cannot be a starting point on this branch. It follows that:

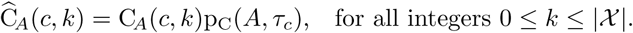

If there is no starting point of continuous *A*-paths in the subtree *𝒯* (*c*) and if no continuous *A*-path passes through *c*, then the same holds for *n* in 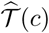We have

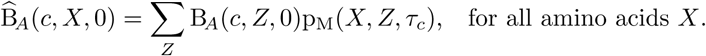

For all positive integers *k*, observing *k* + 1 starting points on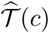 and no *A*-path passing through *n* may come from two mutually exclusive possibilities: either there were already *k* + 1 starting points on *𝒯* (*c*) and no *A*-path passing through *c*, or there were only *k* starting points on *𝒯* (*c*) and an *A*-path passing through *c* which starts on the branch between *n* and *c*. These two possibilities correspond to the terms at the right of the sums below. We have that, for all integers 0 *≤ k ≤ |*𝒳*|*,

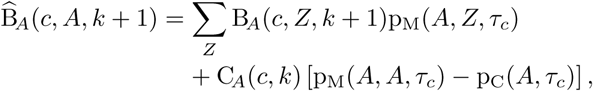

and, for all amino acids Y ≠ A,

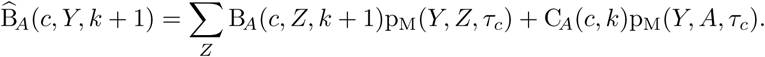

For the sake of simplicity, we assume that *n* has only two children *c*_1_ and *c*_2_. We can actually handle polytomies, but the computations are quite more complicated to write down. Let us make two general remarks:

1. under the model, the evolution on the parts of the trees 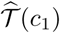 and 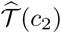 is independent conditionally on the amino acid present at *n*;
2. the total number of starting points on *𝒯* (*n*) is the sum of that on 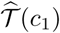 and of that on 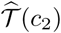.

Since an *A*-path passing through *n* may come either from *c*_1_ or from *c*_2_ or from both of them, we get that, for all integers 0 *≤ k ≤ | 𝒳 |*,

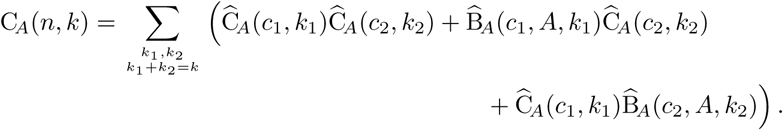

There is no *A*-path passing through *n* if and only if there is both no *A*-path reaching *n* from *c*_1_ and no *A*-path reaching *n* from *c*_2_. It follows that, for all integers 0 *≤k ≤| 𝒳 |*,

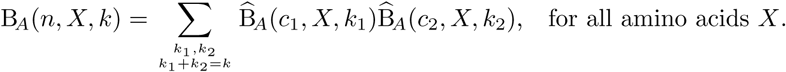

By convention, if there is an *A*-path which starts before the root *r* of *𝒳*, we count an extra starting point. The probability of observing *k* starting points on 𝒳 with an *A*-path passing through *r*, thus counting for *k* + 1, is the product of the conditional probability C_*A*_(*r, k*) with *π*_*A*_, i.e. the probability of amino acid *A* in the initial distribution of the evolution model. The probability of observing *k* starting points on *𝒳* with no *A*-path passing through *r* is the sum over all the amino acids *Z*, of the product of this probability conditioned on having *Z* at the root, i.e. B_*A*_(*r, Z, k*), with the initial probability *π*_*Z*_. We get that the *A*-expectation is:

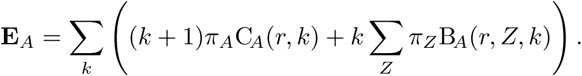

Finally, by putting P({α(𝓁)}_𝓁∈ℒ_) for the probability of observing the amino acids of the extant species under the evolution model (*π, Q*) (i.e. the probability of the tip configuration computed using the pruning algorithm of Felsenstein (1981)), we define the *convergence index* of the site as

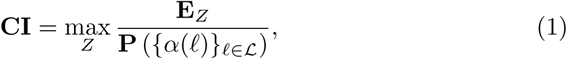

that is the maximum over all the amino acids *Z*, of the expected number of independent apparitions of *Z* going to an extant species with the convergent trait, conditioned on the amino acids of the extant species.

